# MICRORNAS AS REGULATORS OF DRUG METABOLISM AND TRANSPORT IN PREGNANT AND LACTATING WOMEN

**DOI:** 10.1101/2025.07.18.665636

**Authors:** R Fichorova, J Dreyfuss, H Pan, S Nartey, H Yamamoto, P Chen, X Gao, GF Doncel, R Barbieri

## Abstract

**Background:** Physiological changes during pregnancy result in altered maternal drug metabolism that impacts efficacy and safety of therapeutics in pregnant women. Pregnancy induced hormonal, immunologic, or metabolic changes may also influence and alter drug disposition. Despite research efforts focused on pharmacokinetics of medications used in pregnant women in the past decade, knowledge gaps exist in understanding how pregnancy influences drug disposition and placental drug transporters. Moreover, there is a scarcity of research in understanding the safety and effectiveness of therapeutics in lactating women. This study aimed to determine the effect of pregnancy on levels of miRNAs regulating drug metabolizing enzymes and transporters (DMET).

**Methods:** We utilized longitudinal serum specimens collected in 3-month intervals from 88 women who became pregnant during follow-up in a large prospective study of hormonal contraception and HIV acquisition in Uganda and Zimbabwe. We used the HTG EdgeSeq platform coupled with Illumina sequencing to obtain the global miRNA transcriptome in paired specimens collected before, during and after pregnancy. To identify differentially expressed (DE) miRNAs that distinguish pregnancy from pre-conception or breastfeeding we used mixed effect model accounting for multiple samples from the pregnancy event and controlling for fixed effects of batch, country, Nugent score category and sexually transmitted infections. To identify hormonally regulated miRNAs independently associated with Box-Cox-transformed levels of progesterone (P4), β-estradiol (E2), and sex-hormone binding protein (SHBG) we controlled in addition for age, pregnancy and breastfeeding P-values were corrected using the Benjamini-Hochberg false discovery rate (FDR). DMET-targeting miRNAs were identified using miRTarBase focusing on interactions verified by 3’-UTR luciferase reporter assay and overlapped with DE miRNAs with FDR < 0.05.

**Results:** Of 140 DMET-targeting miRNAs among the 2079 miRNAs in the global peripheral blood transcriptome, 41 unique DMET-targeting miRNAs were found to be DE during pregnancy – 38 differentiating pregnancy from preconception and 9 differentiating pregnancy from breastfeeding. The 56 DMETs confirmed as targets of the DE miRNAs included 8 members of the ABC (ATP-binding cassette) transporter family, all abundantly expressed in the placenta, and 4 members of the cytochrome P450 Phase 1 enzyme family with major role in xenobiotics detoxification. The study also revealed a strong (FDR<0.05), predominantly positive association between specific DMET-targeting miRNAs and sex hormone-binding globulin (SHBG) levels, suggesting a miRNA-mediated downregulation of DMETs as SHBG levels rise during pregnancy.

**Conclusion:** This research provides crucial insights into the molecular mechanisms underlying altered drug disposition in pregnant and lactating women, paving the way for improved therapeutic management and personalized medicine in these populations.

## INTRODUCTION

Due to physiological changes during pregnancy, maternal drug metabolism is altered, which affects the efficacy and safety of therapeutics ^1^. Changes in hormones, immunity, and metabolism that are caused by pregnancy can also influence molecular mechanisms underlying drug disposition ^2^. The past decade has seen significant research efforts focused on medication pharmacokinetics in pregnant women, but knowledge gaps persist regarding how pregnancy affects drug disposition and placental drug transporters ^3^. A lack of research is also present in understanding the safety and effectiveness of therapeutics during lactation. In this study, we investigated how pregnancy and breastfeeding affect levels of micro(mi)RNAs regulating drug metabolizing enzymes and transporters (DMETs). MiRNAs are non-coding RNAs of 20-24 nucleotides linking epigenetic transcriptional to post-transcriptional gene regulation by inhibiting translation or degrading mRNA thus controlling protein levels including those of DMETS, mostly explored in relevance to cancer^4 5^. This study fills in the gap by investigating the global miRNA peripheral blood transcriptome changes associated with pregnancy and lactation. We hypothesized that a) miRNA validated experimentally and through predicted sequence-specific base-pairing with DMET mRNAs will be differentially expressed (DE) during pregnancy compared to preconception or breastfeeding, and b) a specific set of DMET-targeting miRNAs will be hormonally regulated.

## MATERIALS AND METHODS

### Study participants

This study included longitudinal samples from 88 women who became pregnant during a large prospective study designed to study hormonal influences on immunity and susceptibility to infection in Uganda and Zimbabwe. The study enrolled HIV-negative women aged 18-35 years (mean age of the 88 pregnant women 23.5 +/-3.5) and followed them with study visits every 12 weeks for a median of 21.5 months. Serum samples were collected from all participants in three-month intervals, allowing analysis of non-pregnant/preconception visit, the next quarterly visit when they were pregnant and the first visit available post-partum when they were non-pregnant and breastfeeding ^6-9^. The study collected information on country, age, sexually transmitted infections (including HIV seroconversion, genital herpes, chlamydia, gonorrhea, candidiasis, trichomoniasis, syphilis) and vaginal dysbiosis assessed by Nugent scoring into categories of bacterial vaginosis, intermediate microflora and normal microflora. There were a few missing values in candidiasis and trichomoniasis infection status, and Nugent score category. These were imputed from the previous visit, unless the missing value was from the first visit, in which case it was imputed from the following visit.

### Global transcriptome analysis of circulating miRNAs

We used the HTG EdgeSeq platform^10^ coupled with Illumina sequencing as described in detail elsewhere to obtain the global miRNA transcriptome in paired specimens collected before, during and after pregnancy.

miRNAs with low counts were filtered out, counts were voom-transformed to log2 counts per million^11^ and normalized by using the voomWithDreamWeights function of R package variancePartition^12,13^. To get an overall view of the similarity and/or difference of the samples, we performed principal component analysis (PCA) using the prcomp function of R package stats R version 4.3.1 (2023-06-16 ucrt).

### Differential expression analysis

We used R version 4.3.1 (2023-06-16 ucrt) and linear mixed effect models to identify differentially expressed miRNAs that distinguish pregnancy from preconception or from breastfeeding following pregnancy by using the dream function of R package variancePartition ^12,13^. The mixed effect models incorporated the feature of multiple specimens from the same pregnancy event (i.e., participant as the random effect) and controlled the fixed effects of sample batch (HTG plate), country, Nugent score category (low, medium, and high), and sexually transmitted infection (including HIV, Candidiasis, Chlamydia, Gonorrhea, HSV, and TV infection status). Further, we computed moderated t-statistics of differential expression by using the eBayes function which implemented empirical Bayes moderation of the standard errors towards a global mean standard error ^14^. Supplemental Table 1 provides average transformed levels of expression for the prepregnant, pregnant and breastfeeding visits, and p values, false discovery rate (FDR) and log Fold Change (FC) from each comparison for each of the 2079 miRNA that showed more than 4 counts per million (CPM) in more than 10 biospecimens.

### Assessment of hormonal levels and their association with miRNA levels

Systemic levels of endogenous β-estradiol (E2), progesterone (P4) were quantified using an Meso Scale Discovery (MSD) electrochemiluminescence duplex assay (MSD, Gaithersburg, MD) and sex-hormone binding globulin (SHBG) levels were measured by Luminex assay (R&D Syetms-Bio-Techne, Minneapolis, MN).

We used mixed effect models to identify hormonally regulated miRNAs, which were associated with Box-Cox-transformed levels of E2, P4, the E2/P4 ratio and SHBG. As applied for DE miRNA above, the mixed effect models incorporated participant as the random effect to account for multiple measurements in each single pregnancy event, and controlled the fixed effects of batch, sample group (preconception, pregnant, and breastfeeding), country, age, Nugent score category, and sexually transmitted infection. P-values were corrected using the Benjamini-Hochberg false discovery rate (FDR) ^15^. P values, FDR and slope from the regression test are provided in Supplemental Table 2.

### Identifying DMET targets

We obtained drug drug-metabolizing enzymes list from INTEDE database ^16^ and drug transporter list from VARIDT database ^17,18^. Also, we obtained the miRNAs and their targeting gene list from miRTarBase ^19^. There were many interactions between miRNA and DMET genes, so we restricted to those that were verified by 3’-UTR luciferase reporter assay and established their overlap with miRNAs significantly up-or down regulated by pregnancy or breastfeeding with FDR < 0.05. The graph showing the interaction of the overlapped miRNA and their DMET gene targets was made by R package igraph.

## RESULTS and DISCUSSION

### DMET-targeting miRNAs differentially expressed during pregnancy and breastfeeding

To get an overall view of the similarity and/or difference of the samples, we performed PCA. The PCA plot is shown in Fig. 1A, visualizing that the samples of pregnant and breastfeeding women were globally different from those of preconception women. Volcano plots (Fig. 1B-C) highlight down and upregulated miRNAs and a heatmap (Fig. 1D) visualizes top miRNAs changes at the transition of preconception/pregnancy to pregnancy and pregnancy to breastfeeding.

**Fig. 1.**
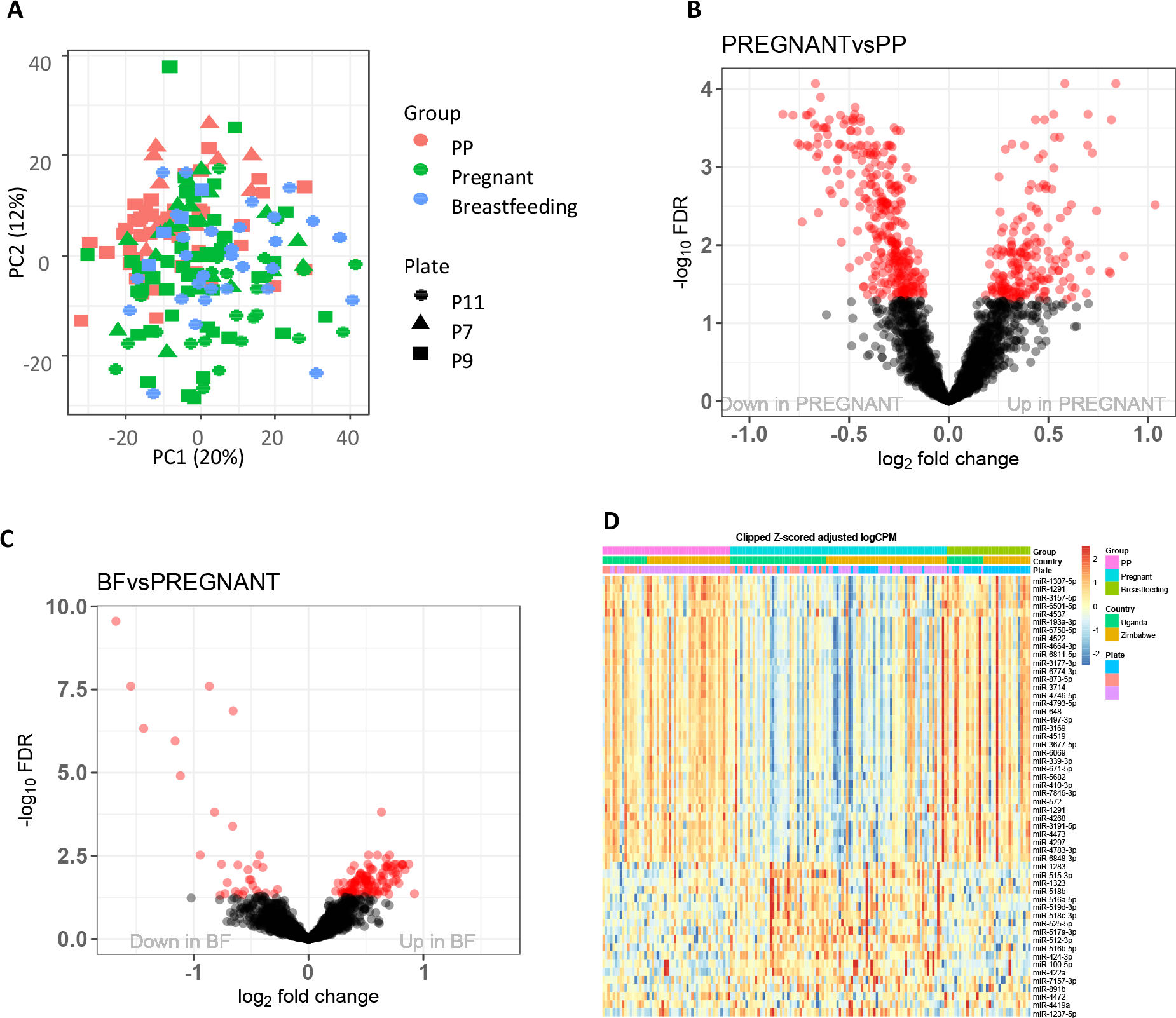
Pregnancy and breastfeeding regulate miRNA expressing. (A) PCA plot using all samples and all miRNA. (B) Volcano plot of pregnancy vs. preconception comparison. (C) Volcano plot of breastfeeding vs. pregnancy comparison. In both (B) and (C), miRNA of FDR < 0.05 are colored in red.(D) Heat map of miRNAs that are significantly changed at the transition from preconception (pregnancy PP) to pregnancy and back to breastfeeding.

The linear mixed model identified 484 miRNAs differentially expressed at the transition from preconception to pregnancy and 147 miRNAs differentially expressed at breastfeeding following pregnancy with an FDR cutoff 0.05 (Supplemental Table 1) The FDR and log fold-change of these significant miRNAs are shown in the volcano plots Fig1. B and C.

Of the 2079 miRNA representing the global miRNAome after filtering out only one miRNA due to low expression, we identified 140 miRNAs which target DMETs according to miRTarBase and selecting interactions confirmed via 3’-UTR luciferase reporter assays. Of these 140 miRNAs, 29% (41 unique miRNAs) were differentially expressed in pregnancy, regulating 56 DMET proteins (Fig. 2 and Tables 1 and 2). Among them, 38 DMET-targeting miRNAs overlapped with miRNAs that differentiated pregnancy and preconception and 9 overlapped with miRNAs that differentiated breastfeeding and pregnancy. These 38 and 9 miRNAs and their DMET targets networks are shown in Fig 2. A and B. Table 1 provides a detailed list of DMET genes targeted by pregnancy-, breastfeeding-, and hormone-regulated miRNAs, directly involved in their disposition, highlighting their class, family, function, and placental expression. It shows 18 experimentally confirmed targets of 20 unique DE miRNAs, representing the major DMET categories of the drug transporter families (ABC and SLC), Phase 1 enzymes involved in oxidation, reduction, or hydrolysis of drugs, Phase 2 enzymes which catalyze their conjugation and members of the nuclear receptor (NR) superfamily directly regulating the transcription of DMETS. All pregnancy DE miRNA-regulated ABC transporters have been shown to be abundantly expressed by the placenta^20-22^. Table 2 presents the remaining DMET-related proteins, their primary function and DE miRNA targeting them. Interestingly, all nine proteins targeted by miRNA differentially expressed by both pregnancy versus preconception and breastfeeding versus pregnancy (mi-873-5p targeting ABCB1, mir-1291 targeting both ABCC1 and SLC2A1, mir299-3p targeting ABCE1 and FXN, miR-518c-3p targeting HSD17B1, mir-193a-3p targeting SLC7A5 and mir-377-3p) followed the pattern returning to pre-pregnancy levels postpartum at the time of breastfeeding (Table 2). Among those mir-518c-3p and another miRNA mir-522-3p (targeting FXN and PTGR1) belong to the C19MC cluster known to be predominantly expressed in the trophoblast^23^, and were decreased significantly (FDR <0.05) postpartum with no significant correlation with SHBG (Table 2). HSD17B1 (enzyme hydroxysteroid 17-beta dehydrogenase 1) is expressed in the placenta, ovaries, endometrium and adipose tissue with primary function of increasing estrogen activity by converting estrone (E1) to estradiol (E2)^24^ and its broader involvement in steroid metabolism makes an important DMET target. Multiple roles have been proposed for FXN (frataxin) particularly in iron metabolism and mitochondrial function and its role in pregnancy is still to be fully elucidated.^25^ PTGR1 (prostaglandin reductase 1) is involved in prostaglandin and other lipid mediators metabolism and in detoxification of ketones and alkenals which are products of oxidative stress naturally increased in pregnancy. While PTGR1 is studied as anti-inflammatory factor in cancer^26^, the decreased levels of the mir522-3p targeting this protein suggest the importance of its natural increase in lactating women which should be further studied.

**Fig. 2.**
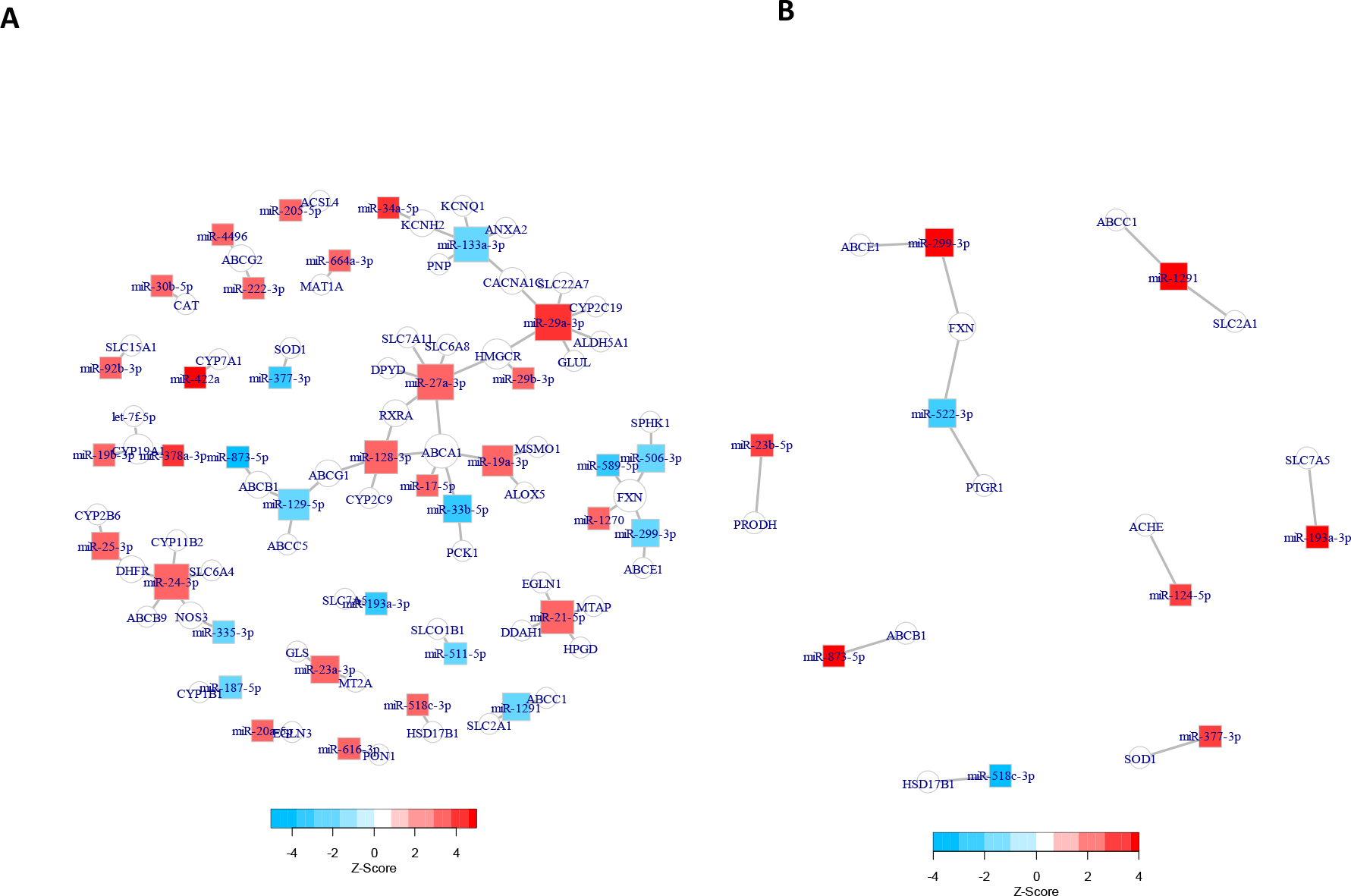
Interaction graph of pregnancy or breastfeeding-regulated miRNAs and their DMENT targets.(A) Pregnancy-regulated miRNAs and their DMENT targets. (B) Breastfeeding-regulated miRNAs and their DMENT targets. In both (A) and (B), the changes of miRNA are indicated by Z-scores (of which the magnitude is determined by *p*-values and sign by log fold-change). The red and blue colors indicate up or down regulation.

**Table 1.**
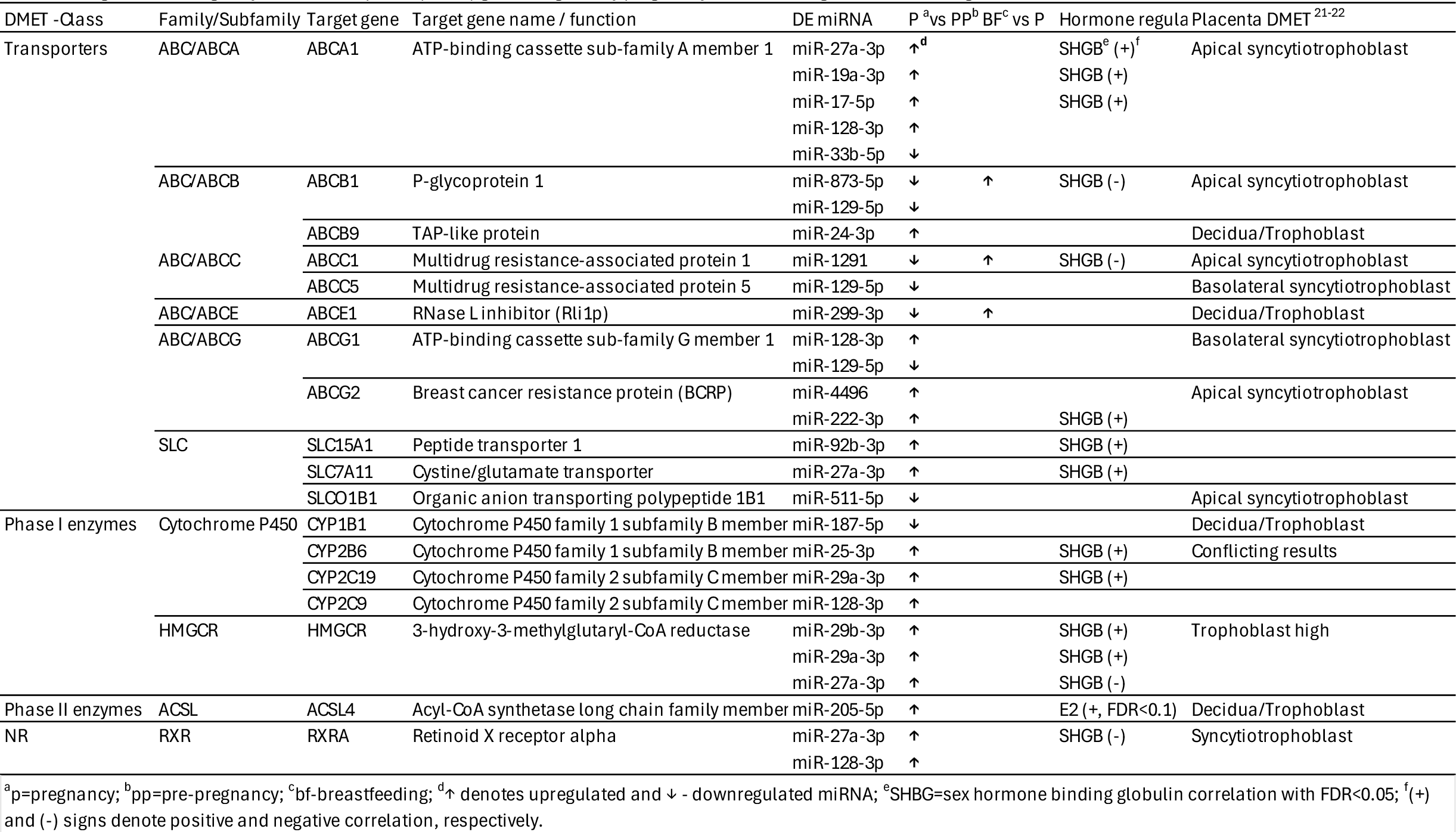
Drug metabolizing enzyme and transporter (DMET) genes targeted by pregnancy, breastfeeding and hormone-regulated miRNA.

**Table 2.** Pregnancy, breastfeeding and hormone-regulated miRNA targeting gene products influencing levels of drug metabolizing enzymes and transporters.

| Target gene | Target gene name / function | DE miRNA | P<sup>a</sup>vs PP<sup>b</sup> | BF<sup>c</sup>vs P | SHBG<sup>d</sup> |
| --- | --- | --- | --- | --- | --- |
| ACHE | Carboxylic ester hydrolase | miR-124-5p |  | ↑<sup>e</sup> |  |
| ALDH5A1 | NAD/NADP acceptor oxidoreductase | miR-29a-3p | ↑ |  | SHGB (+)<sup>f</sup> |
| ALOX5 | Oxygen single donor oxidoreductase | miR-19a-3p | ↑ |  | SHGB (+) |
| ANXA2 | Annexin A2 | miR-133a-3p | ↓ |  |  |
| CACNA1C | Voltage-gated calcium channel alpha Cav1.2 | miR-29a-3p | ↑ |  | SHGB (+) |
|  |  | miR-133a-3p | ↓ |  |  |
| CAT | Peroxidase | miR-30b-5p | ↑ |  | SHGB (+) |
| CYP11B2 | Iron-sulfur protein donor oxidoreductase | miR-24-3p | ↑ |  |  |
| CYP19A1 | Flavin/flavoprotein donor oxidoreductase | miR-378a-3p | ↑ |  |  |
|  |  | miR-19b-3p | ↑ |  | SHGB (+) |
|  |  | let-7f-5p | ↑ |  | SHGB (+) |
| CYP7A1 | Flavin/flavoprotein donor oxidoreductase | miR-422a | ↑ |  |  |
| DDAH1 | Linear amidine hydrolase | miR-21-5p | ↑ |  | SHGB (+) |
| DHFR | NAD/NADP acceptor oxidoreductase | miR-25-3p | ↑ |  | SHGB (+) |
|  |  | miR-24-3p | ↑ |  |  |
| DPYD | NAD/NADP acceptor oxidoreductase | miR-27a-3p | ↑ |  | SHGB (+) |
| EGLN1 | 2-oxoglutarate donor oxidoreductase | miR-21-5p | ↑ |  | SHGB (+) |
| EGLN3 | 2-oxoglutarate donor oxidoreductase | miR-20a-5p | ↑ |  | SHGB (+) |
| FXN | Oxygen acceptor oxidoreductase | miR-1270 | ↑ |  |  |
|  |  | miR-589-5p | ↓ |  | SHGB (-) |
|  |  | miR-506-3p | ↓ |  |  |
|  |  | miR-299-3p | ↓ | ↑ |  |
|  |  | miR-522-3p |  | ↓ |  |
| GLS | Linear amide hydrolase | miR-23a-3p | ↑ |  | SHGB (+) |
| GLUL | Amide synthase | miR-29a-3p | ↑ |  | SHGB (+) |
| HPGD | NAD/NADP oxidoreductase | miR-21-5p | ↑ |  | SHGB (+) |
| HSD17B1 | NAD/NADP oxidoreductase | miR-518c-3p | ↑ | ↓ |  |
| KCNH2 | Voltage-gated potassium channel Kv11.1 | miR-34a-5p | ↑ |  |  |
|  |  | miR-133a-3p | ↓ |  |  |
| KCNQ1 | Voltage-gated potassium channel Kv7.1 | miR-133a-3p | ↓ |  |  |
| MAT1A | Alkyl/aryl transferase | miR-664a-3p | ↑ |  |  |
| MSMO1 | NADH/NADPH donor oxidoreductase | miR-19a-3p | ↑ |  | SHGB (+) |
| MT2A | Quinone acceptor oxidoreductase | miR-23a-3p | ↑ |  | SHGB (+) |
| MTAP | Pentosyltransferase | miR-21-5p | ↑ |  | SHGB (+) |
| NOS3 | NADH/NADPH donor oxidoreductase | miR-24-3p | ↑ |  |  |
|  |  | miR-335-3p | ↓ |  |  |
| PCK1 | Carboxy-lyase | miR-33b-5p | ↓ |  |  |
| PNP | Pentosyltransferase | miR-133a-3p | ↓ |  |  |
| PON1 | Carboxylic ester hydrolase | miR-616-3p | ↑ |  |  |
| PRODH | Quinone acceptor oxidoreductase | miR-23b-5p |  | ↑ |  |
| PTGR1 | NAD/NADP acceptor oxidoreductase | miR-522-3p |  | ↓ |  |
| SLC22A7 | Organic anion transporter 2 | miR-29a-3p | ↑ |  | SHGB (+) |
| SLC2A1 | Glucose transporter type 1 | miR-1291 | ↓ | ↑ | SHGB (-) |
| SLC6A4 | Sodium-dependent serotonin transporter | miR-24-3p | ↑ |  |  |
| SLC6A8 | Creatine transporter 1 | miR-27a-3p | ↑ |  | SHGB (+) |
| SLC7A5 | L-type amino acid transporter 1 | miR-193a-3p | ↓ | ↑ | SHGB (-) |
| SOD1 | Superoxide radical acceptor oxidoreductase | miR-377-3p | ↓ | ↑ |  |
| SPHK1 | Phosphotransferase | miR-506-3p | ↓ |  |  |

<sup>a</sup>p=pregnancy; <sup>b</sup>pp=pre-pregnancy; <sup>c</sup>bf-breastfeeding; <sup>d</sup>SHBG=sex hormone binding globulin correlation with FDR<0.05; <sup>e</sup>↑ denotes upregulated and ↓ - downregulated miRNA; <sup>f</sup>(+) and (-) signs denote positive and negative correlation, respectively.

### Hormonal regulation of DMET-targeting miRNAs

To discover the miRNAs that were associated with hormone E2, P4, SHBG, and E2/P4 ratio, using an FDR cutoff 0.05, we identified 3 miRNAs were significantly associated with E2, while 239 miRNAs were significantly associated with SHBG. Among these miRNAs, 1 was significantly associated with both E2 and SHBG While only 35% of all SHBG-associated miRNAs with FDR <0.05 showed a positive correlation with SHBG levels, enrichment of positive correlation (85%) was shown among the key DMET transporters and metabolizing enzymes shown in Table 1 (11/19 unique miRNA targeting the DMETS were positively correlated with SHBG). The remainder of the 41 DEmiRNA showed the same prevalent positive association with SHBG (Table 2). These relationships suggested a prevailing miRNA-mediated downregulation of key pregnancy DMETS to be expected with rising levels of SHBG observable in the first 24 gestational weeks ^27^. One miRNA (miR-205-5p) targeting ACSL4 (Phase II enzyme) showed a positive correlation with E2 (FDR<0.1). The relationships of all 41 DMET-targeting and pregnancy-regulated miRNAs with the hormones are visualized by heatmaps (Fig. 3A-D).

**Fig. 3.**
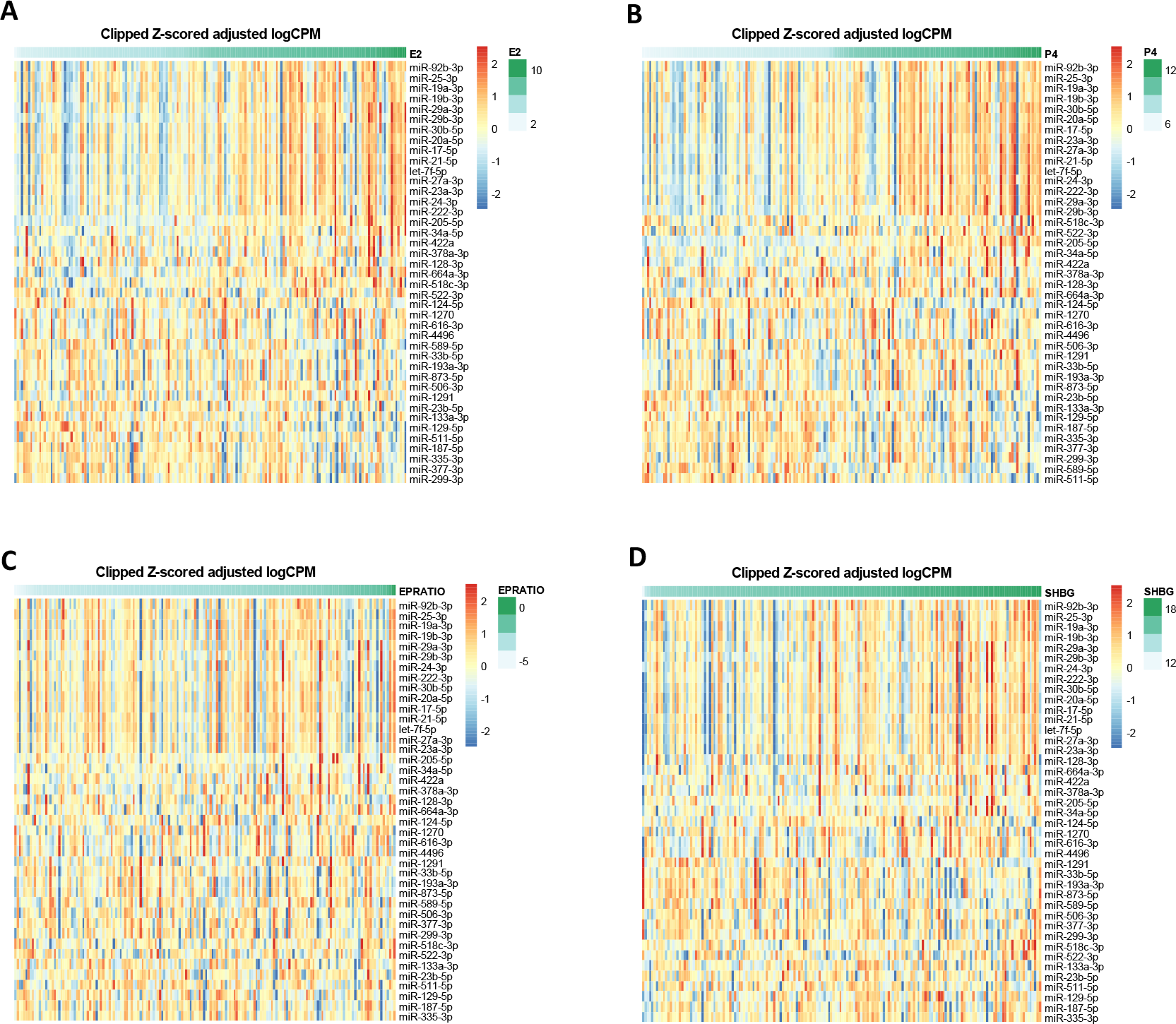
Association of DMET-targeting and pregnancy-regulated miRNAs with the hormones. (A)-(D) Heatmaps show the expression the DMET-targeting and pregnancy-regulated miRNAs that are ordered by hormone expression.

## CONCLUSIONS

This study revealed the profound impact of pregnancy and lactation on maternal drug metabolism and transport, focusing on the regulatory role of microRNAs (miRNAs) on drug metabolizing enzymes and transporters (DMETs). It successfully identified a significant number of circulating miRNAs that are differentially expressed during pregnancy and breastfeeding and regulate crucial DMETs. The findings have critical implications for personalized medicine in pregnancy as identifying specific miRNA signatures could help predict individual variations in drug metabolism and response. A better understanding of miRNA-mediated drug disposition can lead to safer and more effective drug dosing strategies for pregnant and lactating women. The strong correlation between these miRNAs and SHBG levels highlights a novel mechanism for the physiological regulation of drug disposition during pregnancy. The study lays the groundwork for further investigation into the precise mechanisms by which hormones and miRNAs interact to influence drug pharmacokinetics in these vulnerable populations.

